# Phylogenetic analyses reveal an increase in the speciation rate of raphid pennate diatoms in the Cretaceous

**DOI:** 10.1101/104612

**Authors:** Alexandra Castro-Bugallo, Danny Rojas, Sara Rocha, Pedro Cermeño

## Abstract

The raphid pennates (order Bacillariales) are a diverse group of diatoms easily recognized by having a slit in the siliceous cell wall, called the raphe, with functions in cell motility. It has been hypothesized that this morphological innovation contributed to the evolutionary success of this relatively young but species-rich group of diatoms. However, owing to the incompleteness of the fossil record this hypothesis remains untested. Using the 18S ribosomal RNA gene, fossil calibrations, and Bayesian phylogenetic and diversification frameworks, we detect a shift in the speciation rate of marine raphid pennate diatoms in the Cretaceous, not detected in other diatom lineages nor previously recognized in the microfossil record. Our results suggest a positive link between the speciation of raphid pennate diatoms and the benefits derived from evolving motility skills, which could account for their outstanding present-day global diversity. The coincidence between the advent of the raphe and the increase in the speciation rate of raphid pennates supports the idea that simple morphological novelties can have important consequences on the evolutionary history of eukaryotic microorganisms.

## Introduction

Diatoms are silica-precipitating microalgae responsible for roughly one fifth of present day global primary production (i.e. ~20 Gt C per year), and contribute disproportionately to the maintenance of upper trophic levels and carbon sequestration through organic burial in sediments^1,2^. Diatoms are characterized by a peculiar diplontic life cycle involving gradual size reduction during asexual divisions followed by necessary size restitution via sexual reproduction^3,4^. Unlike the centric diatoms, which release flagellated microgametes to the extracellular medium, pennate diatoms normally produce non-flagellated amoeboid gametes of equal size in response to the pairing of vegetative cells of the opposite mating type. Another fundamental difference between centric and pennate diatoms lies in the mating system. Whereas the centrics are strictly homothallic and thus a single clone can produce both male and female compatible gametes, the pennates tend to be heterothallic, that is cross-fertilization must proceed from gametes produced by different clones or mating types^5–7^. The need for size restitution via sexual reproduction has confined pennate diatoms primarily into shallow benthic habitats, where sexual encounters among mating cells become more likely. Still some pennates have adapted successfully to planktonic habitats by maximizing sexual encounters during bloom events. The system of partner location works particularly well in raphid pennates, a lineage of pennate diatoms within the order Bacillariales, easily recognized by having a slit in the siliceous cell wall or frustule, the raphe, which confers the cells autonomous motility^8,9^. Indeed, in spite of their relatively short evolutionary history (i.e. raphid pennates are not known in the fossil record before the Palaeocene epoch), they are the most species-rich group of diatoms today^9,10^, which leads to the hypothesis that this morphological feature facilitated their diversification.

The dynamics of diatom diversity through time has been usually investigated from the analysis of the microfossil record^11–16^. Two major geological projects, the Ocean Drilling Program (ODP) and the Deep Sea Drilling Project (DSDP), now integrated into the International Ocean Discovery Program (IODP) have been instrumental to add new fossil data and unify taxonomic criteria into a global microfossil database with enormous potential for paleoecological research^17,18^. However, the fossil record of marine diatoms is incomplete and severely biased toward recent times for several reasons^13,19–21^. First, silica structures are prone to dissolution at early stages of diagenesis and only a minor percentage of the silica frustules that accumulate in the sediments become eventually preserved. Second, silica recrystallizes under pressure, and as a consequence early diatoms are only preserved through unusual processes such as early carbonate cementation, pyritization or shallow burial. Third, sampling probability decreases with increasing geologic age owing to a deeper position of ancient sediments and the loss of oceanic crust at subduction zones. As a consequence, detailed taxonomic catalogues of fossil diatom assemblages are limited to unconsolidated sedimentary deposits dating back as much as the mid Cenozoic (~40 million years ago, Ma) and thus the evolutionary rates of marine diatoms prior to this time are virtually unknown^20,21^. These preservation biases call into question the suitability of the marine microfossil record for studying the macroevolutionary patterns of eukaryotic microorganisms^9,19^.

It is possible to estimate evolutionary rates (speciation and extinction) through time from molecular phylogenies based on extant species^22^. Distributions of time-calibrated phylogenetic trees are used to detect shifts in the speciation (and extinction) rate through time by simulating distinct evolutionary patterns and searching for the one(s) that best explain the dynamics of the clades within the phylogeny^23–25^. Here we infer a time-calibrated marine diatom phylogeny including available sequences of the 18S ribosomal RNA gene in GenBank and use a reversible jump Markov Chain Monte Carlo method^26^ to detect speciation rate shifts and estimate rate values in major diatom lineages. The method takes into account biases associated with incomplete taxon sampling by incorporating missing lineages at the tree inference stage once provided sampling probabilities. Our objective is to explore the timing and patterns of marine diatom speciation, test for differences among groups, and identify potential causes for their ecological success in the modern oceans.

## Results

### Sequence alignment and phylogenetic analyses

An initial maximum length (without gaps) of 1645 base pairs (bp) resulted in a final alignment of 1157 bp after trimming for terminal regions. We detected that the trimmed sequences of *Thalassiosira pacifica*, *Nitzschia closterium*, *Chaetoceros neogracile*, *Porosira pseudodenticulata* and *Attheya septentrionalis* were equal to *Thalassiosira aestivalis*, *Cylindrotheca fusiformis*, *Chaetoceros gracilis*, *Porosira glacialis* and *Attheya longicornis,* respectively, so the first four sequences were deleted at the haplotypes collapsing step.

Maximum-likelihood (ML) and Bayesian inferences (BI) resulted in similar estimates of the phylogenetic tree, with congruent well-supported relationships (Figure S1, Figure S2). In these "unconstrained" ML/BI searches, most relevant groups obtained high support in accordance with previous phylogenetic analyses^27,28^. Rhizosoleniales, Coscinodiscales, Chaetocerotales, Thalasiosirales and Bacillariales (except *Attheya longicornis*) were highly supported both in Bayesian inference (Posterior probability, PP = 1) and ML (Bootstrap support, BS > 900). The sister-relationship between *Attheya longicornis* and the other Bacillariales included in the analyses was the lowest supported relationship (PP = 0.79; BS =540), but this is in agreement with previous studies showing the uncertain position of *Attheya longicornis* within diatoms ^29^. The remaining Bacillariales received maximum support for defining a monophyletic clade.

Using the calibration points (Table S2), divergence time analyses estimated the root of the group at 193.32 Ma with a 95% highest posterior density (HPD = 121.98-274.74 Ma). This result is also consistent with other phylogenetic studies, which situate the origin of diatoms between the Triassic and lower Jurassic^28,30^. The mean age of the Thalassiosirales clade was 113 Ma (HPD = 69-155), and both Coscinodiscales and Bacillariales (including *Attheya longicornis*) were estimated to have originated in the Cretaceous, 115 Ma (HPD = 62-168) and 118 Ma (HPD = 62-180), respectively. Chaetocerotales and Rhizosoleniales seem to have a more recent origin, ~66 Ma (HPD = 56-86) and 91.5 Ma (HPD = 88-94), respectively (Figure 1, see also information available through the following Figshare DOI: 10.6084/m9.figshare.3795834).

**Figure 1.**
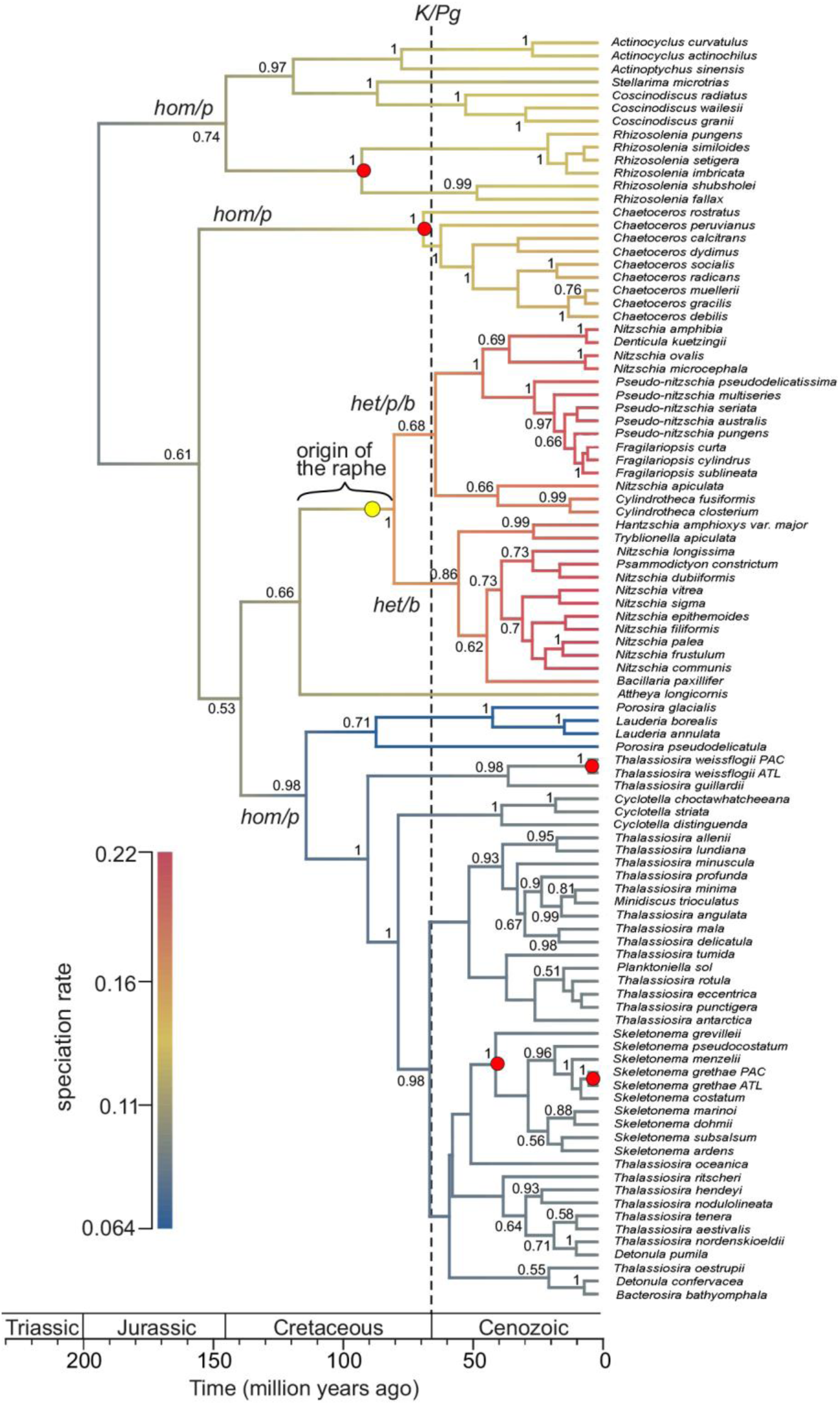
Phylorate plot of the mean speciation rates of marine diatoms sampled from the posterior resulting from the BAMM output. Branches coloring (blue to red) indicates increasing speciation rates. The yellow dot marks the position of the speciation rate shift resulting from the overall best shift configuration. The red dots indicate the calibration points based on the sedimentary record (Table S2). Numbers at branches represent posterior probabilities (only values >0.5 are shown). The abbreviations over major clade branches denote the predominant sexual strategy and lifestyle within lineages: homothallism (hom), heterothallism (het), planktonic (p) and benthic (b). Also indicated is the time span for the presumed origin of the raphe.

### Diversification analyses

The effective sample size for the number of shifts and log-likelihood was superior to 2000 in both cases, indicating the appropriate sampling of parameters from the posterior. The 95% of the most credible rate shift sets were sampled with the Bayesian Analysis of Macroevolutionary Mixture (BAMM) from which twelve shift configurations were generated (see Material and Methods, Table S3). We found strong evidence for the one rate shift macroevolutionary model compared with the zero rate shift model (BF = 19.95). The rate shift was positioned at the base of the raphid pennate diatoms core within the order Bacillariales ~78-80 Ma (Figure 1). The two rate shift macroevolutionary model (BF = 5.03) was discarded because the overall best shift configuration positioned the second shift in the same location as the first one with less significant statistical support (Figure S3, Table S3). This speciation rate shift is the first one detected with non-fossil techniques and is in agreement with the expansion of raphid pennate diatoms during the Cenozoic era.

Phylorate plots showed that the maximum speciation rate of Bacillariales was as high as 0.2 species My^−1^, while the maximum for other diatom groups such as Thalassiosirales never exceeded 0.1 species My^−1^ (Figure 2). All groups exhibited an increasing speciation rate through time. We found statistically significant differences in speciation rates between raphid pennate diatoms and centric diatoms (*F* = 191.96, *P* = 0.0097, significance level = 0.05). The low speciation rate of Thalassiosirales throughout the Cenozoic era is conspicuous because this order is particularly successful in the modern oceans.

**Figure 2.**
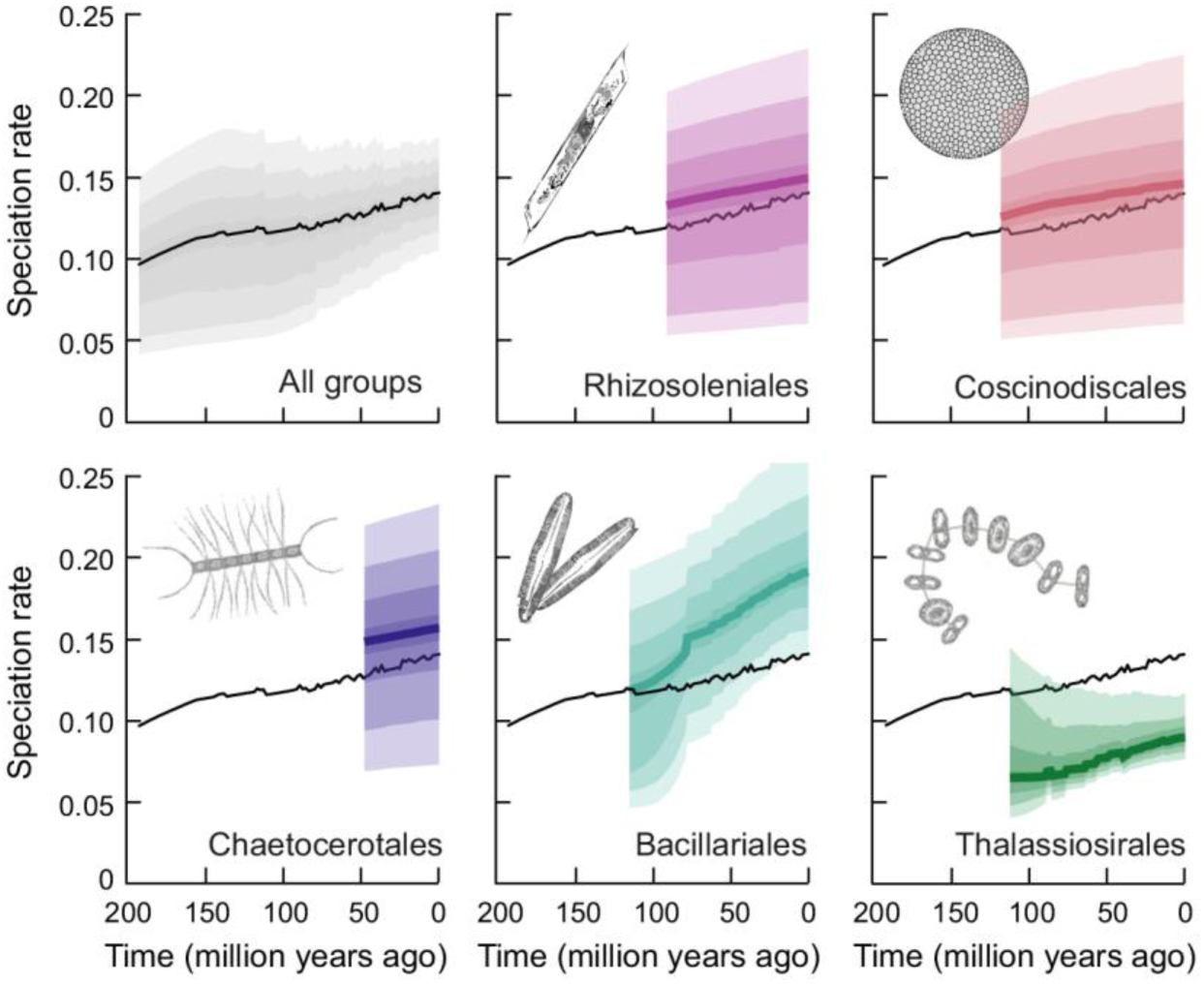
Speciation rate through time plot of the different groups of marine diatoms compared with the evolutionary rate of the entire phylogeny (black line). Bacillariales are the primary responsible for the speciation rate shift detected in the phylogeny. Bacillariales plot includes all the species of the order except *Attheya longicornis*. Colored areas represent the quantiles of the posterior distribution. Pictures were taken by Isabel Gomes Teixeira.

## Discussion

Using time-calibrated diatom phylogenies within a Bayesian framework we detected a shift in the speciation rate of raphid pennates 78-80 Ma. This speciation rate shift could not be previously recognized in the marine diatom fossil record, which is sparse for that time and strongly biased towards recent times^13,18^.

The fossil record situates the origin of raphid pennate diatoms some 70.6-55.8 Ma^19^, yet, our results (Figure 1) and previous molecular analyses push back their origin earlier in the Cretaceous. Analyses of the18S rRNA and ribulose bisphosphate carboxylase/oxygenase large chain genes have shown that the raphid pennates comprise a monophyletic group of diatoms within the order Bacillariales^27,28,30^. Regarding to *Attheya longicornis*, its phylogenetic position is largely uncertain^29^, and although phylogenetic analyses seem to indicate that it shares an ancestor with the raphid pennates, it is clearly a distant relative and lacks a raphe. The inferred speciation rate shift was located in the Bacillariales clade, excluding *Attheya longicornis*, which further suggests that the most probable shift detected in our phylogeny corresponds to the evolutionary expansion of the raphid pennates.

The speciation rate shift reported here could result from i) a greater chance of sexual reproduction in raphid pennate diatoms with respect to other diatom lineages and ii) a tendency towards heterothallism in the pennate diatoms mating system, which may have contributed to erect hybridization barriers and reduce gene flow among sympatric populations.

Meiotic recombination in sexually-reproducing eukaryotes represents a first order cause of genotypic variability within mating populations and hence the frequency of sexual reproduction potentially accelerates the rate of species evolution^31,32^. Sexual reproduction is an obligate stage in the life cycle of most diatoms. However, the success of sexual reproduction may differ among diatom lineages depending on their sexual strategy and lifestyle^9,33^. For instance, in centric diatoms, which have adopted primarily a planktonic lifestyle, environmental cues induce the formation of eggs and flagellated sperm, which is released to the extracellular medium. A suite of concurrent requirements including appropriate environmental cues, the occurrence of sexualized cells and bloom-forming conditions that increase the chance of egg fertilization must be met to succeed^34,35^. Conversely, in pennate diatoms, the production of gametes is preceded by the pairing of vegetative cells of the opposite mating type^33,36^, a process that seems to be facilitated by the release of regulatory pheromones^37,38^. The system of partner location works especially well in raphid pennates, which use their motility skills to search actively for a partner and join their gametes^9^. Though there is little information regarding the frequency of sex in diatoms, previous reports indicate that, overall, pennate diatoms tend to exhibit shorter temporal lags between sexual events than centric diatoms^7^. For instance, the cell size threshold for inducing sexual reproduction in the diatom *Pseudo-nitzschia multiseries* has been shown to be larger than previously thought^39^, expanding the window of opportunity for sexualization.

Sexual reproduction favors adaptation to new habitats and is particularly advantageous in rapidly changing environments, where some genetic variants might be wiped out by new conditions while others might be better adapted and thrive. In addition to increasing the chance of sexual recombination, gliding motility potentially increases resource use efficiency by conferring individuals the ability to seek for optimal light environments and nutrient-rich conditions^40–42^. We recognize that the outstanding current diversity of raphid pennate diatoms could be explained in part as a result of a finer niche differentiation in benthic habitats compared to more homogeneous planktonic ecosystems. However, this argument on its own is unable to explain the lower diversity of araphid pennates today despite their primarily benthic lifestyles and comparable evolutionary origins. Furthermore, our analysis shows that the lineages of raphid pennate diatoms adapted to planktonic lifestyles (e.g. *Pseudo-nitzschia*, *Fragilaria*) also attained higher speciation rates than other planktonic lineages such as the Thalasiosirales, Chaetocerotales, Rhizosoleniales and Coscinodiscales, supporting the idea that the benefits derived from evolving motility skills (the raphe) played a role. The timing of the raphid pennate diatoms speciation rate shift suggests that changes in the extent of continental flooding along with concurrent increases in continental nutrient weathering fluxes^43^ and sedimentary organic matter recycling^44^ provided ideal conditions for habitat expansion since the mid-Cretaceous^19^.

The second critical control on evolutionary tempo deals with the fact that pennate diatoms show a tendency towards heterothallism^35^. It has been suggested that owing to their broad dispersal ranges and astronomical population numbers, microbial species cannot be geographically isolated^45^. Because geographic isolation is a necessary component of allopatric speciation models, global dispersal and the ensuing continuity of gene flow among sympatric populations are thought to lower the rate of speciation^45^. Ubiquitous dispersal is feasible in asexually reproducing microorganisms as long as global dispersal times do not exceed the pace of genetic variability. However, the mating system in heterothallic species is a fundamental control of syngamy, which increases the chance of geographic isolation and promotes the spatial structuring of genetic populations^46^. Our results support the idea that the advent of the raphe, a simple morphological feature involved in cell motility, facilitated sexual encounters among compatible mating types. The greater success of sexual reproduction in predominantly heterothallic taxa led to an increase in the speciation rate of raphid pennate diatoms. These results provide a feasible explanation for the outstanding current diversity of raphid pennates despite their relatively recent origin, and suggest that simple morphological novelties can have important consequences on the evolutionary history of eukaryotic microorganisms. While the signal seems strong in our data, it is important to recognize the limitations of single-gene-phylogenies,^47,48^ and would be extremely valuable if future studies could employ more comprehensive sets of orthologs ^27,49,50^, as these expand in databases, or their sequencing becomes more feasible, to keep testing hypotheses concerning timing and rates of diversification.

## Materials and Methods

### Sequences and alignment

All available sequences (102 sequences) of the 18S ribosomal RNA gene of major marine diatom orders (Thalassiosirales, Chaetocerotales, Rhizosoleniales, Coscinodiscales and Bacillariales) were downloaded from GenBank (Table S1). Sequences were aligned using MAFFT v7.058b^51^ and the G-INS-I algorithm^52^, trimmed using GBlocks automatic parameters^53^ and collapsed into haplotypes using ALTER^54^ resulting in a final alignment with 97 sequences and 1157bp. The minimum number of sequences for defining a conserved position was 50 and 83 for a flanking position. The maximum number of contiguous non-conserved position was 8 while the minimum length of a block allowed after gap cleaning was 10. Alignments and haplotypes information are available through the following Figshare DOI: 10.6084/m9.figshare.3795834.

### Phylogenetic analyses

To determine the best-fit model of sequence evolution for the dataset we used jModelTest v0.1.1 ^55,56^ and the corrected Akaike Information Criterion (AICc)^57^. Maximum-likelihood (ML) and Bayesian (unconstrained) gene phylogenies were estimated using PhyML 3.0^58^ and MrBayes v3.2.1^59^. ML was performed with 100 heuristic searches and its support accessed through 1000 bootstrap replicates. Two Bayesian inference runs of 10 million generations were performed, with default heating parameters, sampled each 1000^th^ sample, and checked for convergence (of node’s PP’s) and congruence (of runs) using AWTY^60^. Both runs were summarized on a 50% majority-rule consensus. All details and outputs are available at DOI: 10.6084/m9.figshare.3795834.

BEAST v2.1.3^61,62^ was used for co-estimating divergence times and (constrained) gene phylogenies using relaxed molecular clock analyses. The nucleotide substitution model was implemented as estimated previously, with parameters to be co-estimated along the run. An uncorrelated lognormal model was used for the clock-rate and posterior estimates were obtained under both the Calibrated Yule and the Birth-Death models for the tree prior. Runs without data were also performed to evaluate prior and joint prior distributions.

Calibrations in the phylogeny were implemented either as minimum or “fixed” (interval) ages. Using data from the marine diatom fossil record, we constrained the minimum ages of several clades in the tree corresponding to the oldest unequivocal fossil belonging to that clade. In all cases these clades were highly supported in the unconstrained phylogenies (PP=1; BS>900/1000). The single exception was *Skeletonema grethae*, where specimens from the Atlantic and Pacific were not inferred as sister-taxa. Because the relationships within the clade they belong were largely unresolved (see Figure S1, S2), with no support either for them not being sister taxa, and the distance between them was very low (<0.3%), similar to the one between the other Atlantic-Pacific pair (*Thalassiosira weissflogii*), and to other Atlantic-Pacific taxa pairs which divergence is assumed to have been initiated at the final closure of the Isthmus of Panama^63^, we constrained this node and still included this calibration in our analyses.

The median age and the interval of probability were obtained from a lognormal distribution of the data (Table S2). Fixed age estimates were used for those clades whose divergence times occurred during a well-dated geologic event. In this case, the median and probability interval were calculated using a normal distribution (Table S2).

Two independent analyses (under each tree prior) were run for 400 million generations, sampling every 400,000. The resulting distributions of parameters were checked for convergence using Tracer v 1.5^64^ ensuring effective sample size values (ESS) to be greater than 200. The posterior distributions of trees were summarized in a maximum clade credibility tree discarding 25% as burnin using median heights for node age estimates.

### Speciation rates and phylANOVA analyses

We used the Bayesian Analysis of Macroevolutionary Mixture (BAMM, www.bamm-project.org) to estimate marginal distributions of speciation and extinction rates for each branch in a phylogenetic tree^65^. BAMM uses reversible jump Markov Chain Monte Carlo (rjMCMC) method to detect automatically rate shifts and sample distinct evolutionary dynamics (speciation and extinction) that best explain the whole diversification dynamics of the clade. The program is designed to work with datasets that contain large numbers of missing species. The method takes into account incomplete taxon sampling in phylogenetic trees by incorporating missing lineages at the tree inference stage once provided clade-specific sampling probabilities^65,66^.

We provided BAMM with the proportion of species sampled per genus (i.e., 1/number of species in genus). Priors were estimated with BAMMTools^26^ using the function “setBAMMpriors”. We set a rjMCMC for 11∙10^6^ generations and sampled every 1,000 generations. We used ESS values to assess the convergence of the run, considering values above 200 as indicative of good convergence. To visualize where in the tree the shifts occurred, we generated mean phylorate plots which represent the mean speciation rate sampled from the posterior at any point in time along any branch of the phylogenetic tree^26^. BAMM identifies a set of most credible rate shifts ordering them by posterior probability. Here, we selected twelve credible rate shift sets based on a Bayes Factor (BF) considering a BF value between 3 and 12 as positive evidence, BF > 12 as strong evidence and BF >150 as very strong evidence^67^. All details and outputs are available at DOI: 10.6084/m9.figshare.3795834.

Phylogenetic analyses of variances were used to test differences in speciation rates among raphid pennate diatoms and centric diatoms using the function phylANOVA in the R-package phytools^68^. phylANOVA compares results based on raw phylogenetic data to those based on phylogenetic model simulations to create an appropriate null distribution^69^.

## Acknowledgments

We thank R. Massana, F.M. Cornejo-Castillo and A. Roura for comments on the manuscript. This research was supported by grants 10PXIB312058PR from Xunta de Galicia and CTM2014- 54926-R from the Spanish Ministry of Economy and Competitiveness. P.C. was supported by Ramon y Cajal contract from the Spanish Ministry.

### Author contribution

ACB and PC conceived and designed the experiment, ACB selected and downloaded the sequences, ACB and SR performed the phylogenetic analysis, ACB and DR did the diversification rate analysis, ACB wrote the first draft of the manuscript and all authors contributed substantially to revisions.

### Competing financial interest

The authors declare no competing financial interest.

